# Translational activity of 80S monosomes varies dramatically across different tissues

**DOI:** 10.1101/2024.06.24.600330

**Authors:** Albert Blandy, Tayah Hopes, Elton J. R. Vasconcelos, Amy Turner, Bulat Fatkhullin, Michaela Agapiou, Juan Fontana, Julie L. Aspden

**Affiliations:** School of Molecular and Cellular Biology, Faculty of Biological Sciences, University of Leeds, LS2 9JT, UK; LeedsOmics, University of Leeds, UK; Astbury Centre for Structural Molecular Biology, University of Leeds, Leeds, LS2 9JT, UK

**Keywords:** 80S monosome, mRNA translation, elongation state, tRNA, cryo-EM, RNA-seq

## Abstract

**Summary:** Translational regulation at the stage of initiation can impact the number of ribosomes translating each mRNA molecule. However, the translational activity of single 80S ribosomes (monosomes) on mRNA is less well understood, even though these 80S monosomes represent the dominant ribosomal complexes i*n vivo*. Here, we used cryo-EM to determine the translational activity of 80S monosomes across different tissues in *Drosophila melanogaster*. We discovered that while head and embryo 80S monosomes are highly translationally active, testis and ovary 80S monosomes are translationally inactive. RNA-Seq analysis of head monosome- and polysome-translated mRNAs, revealed that head 80S monosomes preferentially translate mRNAs with TOP motifs, short 5’-UTRs, short ORFs and are enriched for uORFs. Overall, these findings highlight that regulation of translation initiation, and that protein synthesis is mostly performed by monosomes in head and embryo, whilst polysomes are the main source of protein production in testis and ovary.

**Graphical Abstract:** 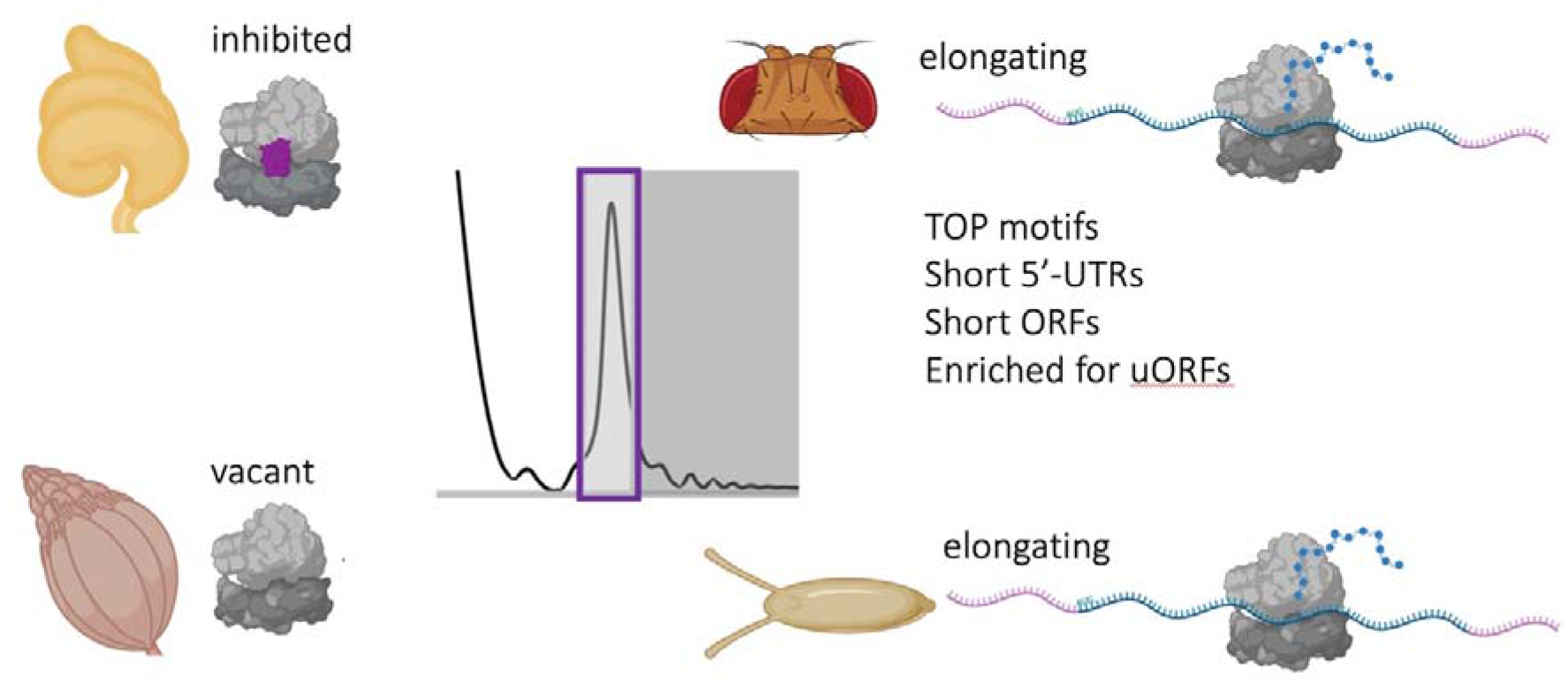

**Highlights:**

- *D. melanogaster* 80S monosomes and polysomes purified from different tissues exhibit different translational activities
- Head and 0-2 hr embryo 80S monosomes are highly translationally active
- Testis and ovary 80S monosomes are translationally inactive
- Head 80S monosomes preferentially translate mRNAs with TOP motifs, short 5’-UTRs, short ORFs and are enriched for uORFs
- Head polysomes preferentially translate mRNAs with neuronal functions

## Introduction

Translation levels are tightly regulated during development and differentiation across eukaryotes (Kong and Lasko 2012). Much of this control acts during the initiation of protein synthesis, regulating the rate of 80S formation and therefore the number of ribosomes bound per mRNA (Hinnebusch, Ivanov, and Sonenberg 2016). mRNAs bound by many ribosomes are termed poly-ribosomes or polysomes (Warner, Knopf, and Rich 1963). The binding of multiple ribosomes on a single mRNA facilitates efficient protein synthesis, therefore there is a generally held assumption is that mRNAs bound by 80S monosomes are less translated than those bound by polysomes. However, several factors complicate this simple assumption, including the length of an ORF and therefore the space available to be bound by multiple ribosomes at the same time (Heyer and Moore 2016; Aspden et al. 2014; Arava et al. 2003).

The proportion of ribosomes engaged as 80S monosomes or polysomes varies across cell types, tissues and response to stimuli (e.g. during differentiation (Douka et al. 2021)). Measuring translation, its regulation and how this differs across cells can be achieved by numerous techniques. The most widely used approach to assess translation levels and how they change, is polysome profiling (Chassé et al. 2017). This involves the separation of ribosomal complexes in sucrose gradients on the basis of the number of ribosomes bound per mRNA, providing an overview of the translational status of a sample. The distribution of individual mRNAs can then be assessed across the gradient to measure what proportion of the mRNA is bound by ribonucleoproteins, 40S subunits, 80S monosomes and polysomes (Douka et al. 2021).

It has been a long-standing assumption that single 80S monosomes consist of 2 types of ribosomes: i) inactive “vacant” 80S, not engaged on mRNAs (Noll et al. 1973); and ii) translationally active 80S, which could be initiating, elongating or terminating. However, the relative proportions of these two types of 80S has not been determined. Dissecting the translational capability of different sized translational complexes, i.e. 80S monosomes vs small polysomes vs large polysomes, has important consequences on understanding rates of protein synthesis and translational regulation. For example, it has previously been shown that in neuronal axons and dendrites, 80S monosomes are highly translationally active and represent a substantial part of the protein synthesis machinery (Biever et al. 2020).

Additionally, ribosome profiling of yeast 80S monosomes has shown that most of them are engaged in elongation, rather than initiation (Heyer and Moore 2016). “Vacant” ribosomes, not engaged in translation, have been shown in several systems to be bound by factors within the mRNA channel preventing mRNA binding. This includes prokaryotic hibernation factors (Helena-Bueno et al. 2024) and eukaryotic dormancy factors, which stabilise 40S and 60S binding to form 80S monosomes, and prevent 40S binding to mRNAs, allowing for regulation during stress or development (Leesch et al. 2023; Brown et al. 2018).

This study set out to determine what proportion of 80S ribosomes are engaged in active translation in *Drosophila melanogaster (*D. melanogaster*)* tissues, and if tissue-specific differences exist. This has revealed a stark difference between translation states of 80S monosomes between tissues. We also found that within heads, 80S monosomes and polysomes translated different mRNA pools, with distinct molecular characteristics and encoding specific protein functions.

Together this work shows that protein synthesis in the head and the embryo is mostly performed by monosomes, in contrast to testis and ovary, where polysomes are the main source of protein production.

## Results

### Cryo-EM of 80S ribosomes from *D. melanogaster* heads at 3.0 Å

To profile the translational activity across different tissues we purified ribosomes from *D. melanogaster* heads, 0-2 hr embryos, testes and ovaries. As we have previously shown, the sucrose gradient profiles from these tissues vary in their precise ratio of 80S monosomes to polysomes (Hopes et al. 2022). However, what is in common between these tissues is that the majority of ribosomes present are 80S monosomes (Figure S1), whilst the minority are polysomes, in contrast to *D. melanogaster S2* tissue culture cells (Hopes et al. 2022; Aspden et al. 2014). Therefore, we sought to understand whether these 80S monosomes were in fact involved in active translation and if this varies across tissues. Cryo-EM was performed on 80S monosomes from heads and a dataset containing ⍰600,000 particles was collected. Image processing of this head 80S dataset as a single class was refined to an average 3.0 Å resolution (Figure S2A).

In contrast to the previous structures that we had determined for testis 80S and ovary 80S monosomes, which contained no tRNA (Hopes et al. 2022), the head 80S map contained tRNAs in A/A and P/P sites, and to a lesser extent the E/E site, which we interpreted as evidence of multiple classes present in the sample. This indicates that the majority of the head 80S monosome are likely engaged in elongation, and therefore actively translating.

### The vast majority of head 80S monosomes contain tRNAs

To dissect the tRNA occupancy of the head 80S monosomes more precisely, we performed focused classification around the mRNA channel and A/P/E tRNA sites of the head 80S 3D average. To ensure potential classes were not limited by the image processing protocol, we attempted classification into 10 classes. This resulted in 6 significant classes (which we defined as containing >1% of the dataset particles and >200 particles) from the head 80S monosome dataset (Figure 1A-F). 5 of 6 of these classes contained tRNAs occupying various sites within the mRNA channel, reflective of different states during translation elongation. Quantification of the number of particles contributing to these classes revealed that ∼61% head 80S monosomes had tRNAs in A/A and P/P sites (Figures 1A-C), ∼19% in P/E and A/P sites (Figure 1D) and ∼19% in P/P alone (Figure 1E). Only ∼2% of the ribosomes had no tRNAs present (Figure 1F).

**Figure 1:**
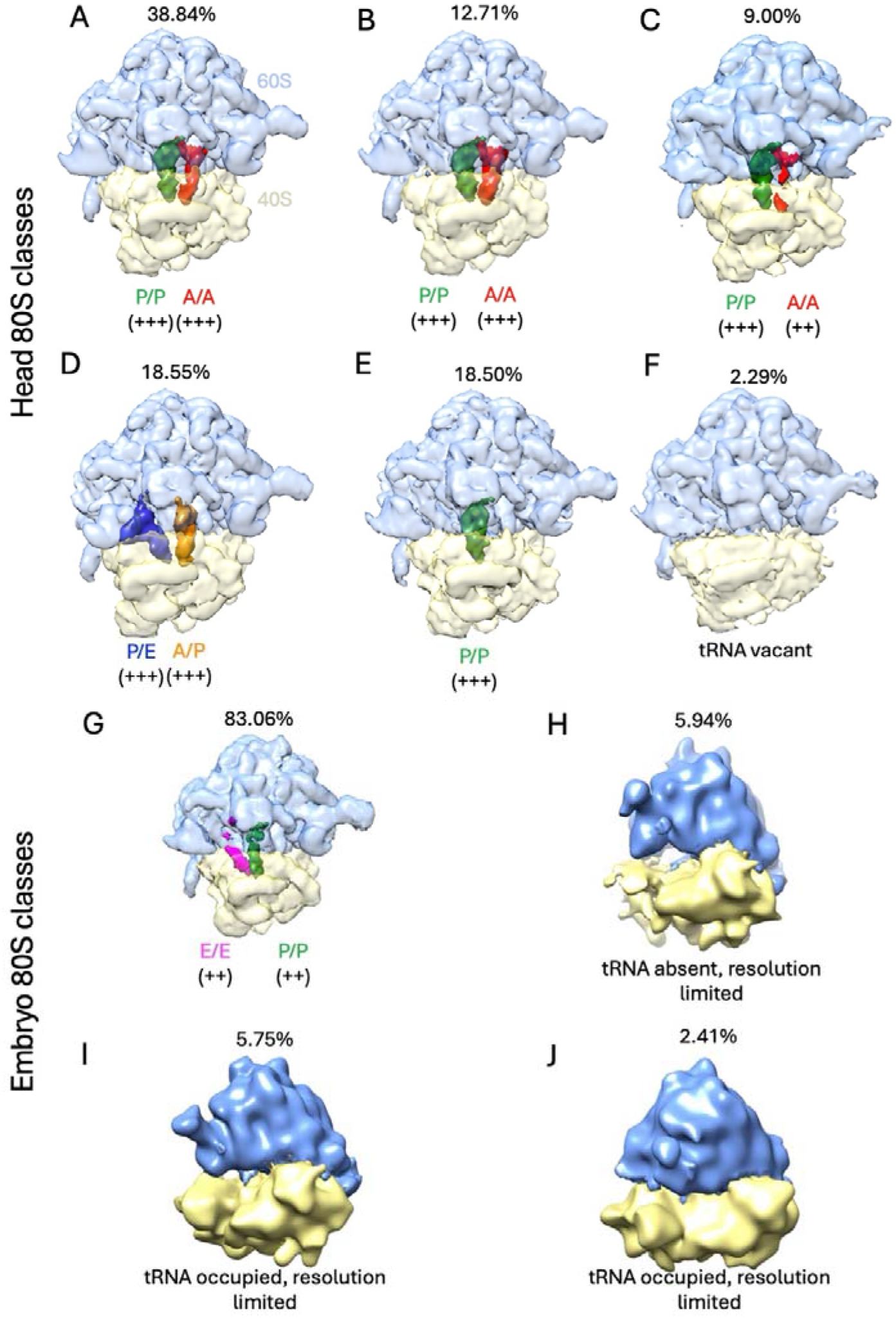
80S monosomes from *D. melanogaster* head and embryo are actively translating. Significant 3D classes (>1% total particles in dataset and >300 particles) of the head 80S dataset (A-F) from ∼600,000 particles; and 0-2 hr embryo 80S monosome dataset (G-J) from ∼11,500 particles. (A to C) 3 classes of the head 80S dataset, representing 38.84%, 12.71% and 9.00% of the head 80S particles, contained tRNA densities in P/P and A/A sites. (D) 18.55% of the head 80S particles (1 class) contained tRNA densities in P/E and A/P sites. (E) 18.50% of head 80S particles (1 class) contained tRNA densities in P/P sites. (F) 2.29% head 80S particles were tRNA vacant. (G) 83.06% of the embryo 80S particles (1 class) contained tRNA densities in E/E and P/P sites. (H) 5.94% of embryo 80S particles (1 class) were tRNA vacant. (I and J) 2 classes, representing 5.75% embryo and 2.41% of embryo 80S particles, contained densities within the mRNA channel and therefore are tRNA occupied, but resolution is limited. Percentage of particles in each class is given above each class. tRNA site positions are indicated along with score of confidence with ++ medium confidence and +++ highly confident.

Additionally, we generated a second cryo-EM head 80S monosomes data set from a biological replicate, which contained ∼7,700 particles. The focused classification of the mRNA channel and tRNA sites of this sample generated two major classes, both with tRNAs present: ∼52% in the A/P and P/E sites and ∼47% in the P/P site (Figure S2B-C). Overall, this analysis indicates that ∼98-99% of 80S monosomes are engaged in active translation in the head.

### Most 80S monosomes from embryo exhibit tRNA occupancy

To profile the translational activity in the same way for embryo 80S monosomes we performed cryo-EM on 80S monosomes purified from 0-2 hr embryos and a dataset containing ⍰11,500 particles was collected. Image processing of this embryo 80S dataset resulted in an average at 5.4 Å resolution. We again performed focused classification around the mRNA channel and tRNA sites, resulting in 4 significant classes. Although fewer particles were contributing to this dataset meant some classes were resolution limited, we could still determine whether they were tRNA occupied or absent (Figure 1G-J).

Like in the head, the vast majority (∼91%) of these embryo 80S monosomes contained tRNA, reflective of active translation elongation (Figure 1G, I, J). The main tRNA occupied state detected in embryo converged into a single class that contained densities for E/E and P/P tRNAs. Together with the head results, this confirms that monosomes can be actively engaged in translation in specific tissues, in contrast to the large abundance of polysomes present in tissue cultured cells.

### No monosomes are engaged in active translation in ovaries or testes

The averages we had previously generated for testis and ovary 80S monosomes did not contain tRNAs (Hopes et al. 2022). These datasets consisted of ∼47,000 and ∼186,000 particles respectively, which resulted in cryo-EM averages at 3.5 and 3.0 Å resolution. However, no focused classification around the mRNA channel/tRNA sites was performed on these datasets. To determine if small proportions of particles did contain tRNAs, and therefore were actively translating, we performed focused classification and quantification on these datasets. We were unable to detect any testis 80S monosomes with tRNAs present. Instead, 2 significant classes were identified, both with the mRNA channel occupied by IFRD1 (Figures 2A-B). Building on our previous work showing the presence of IFRD1 (Hopes et al., 2022), this analysis revealed that all 80S monosomes in the testis are inactive, inhibited by the presence of IFRD1. Additionally, all 4 significant classes of ovary 80S monosomes had empty mRNA channels, with no tRNA density present (Figures 2C-F). Together, this analysis reveals that none of the testis or ovary 80S monosomes are actively translating.

**Figure 2:**
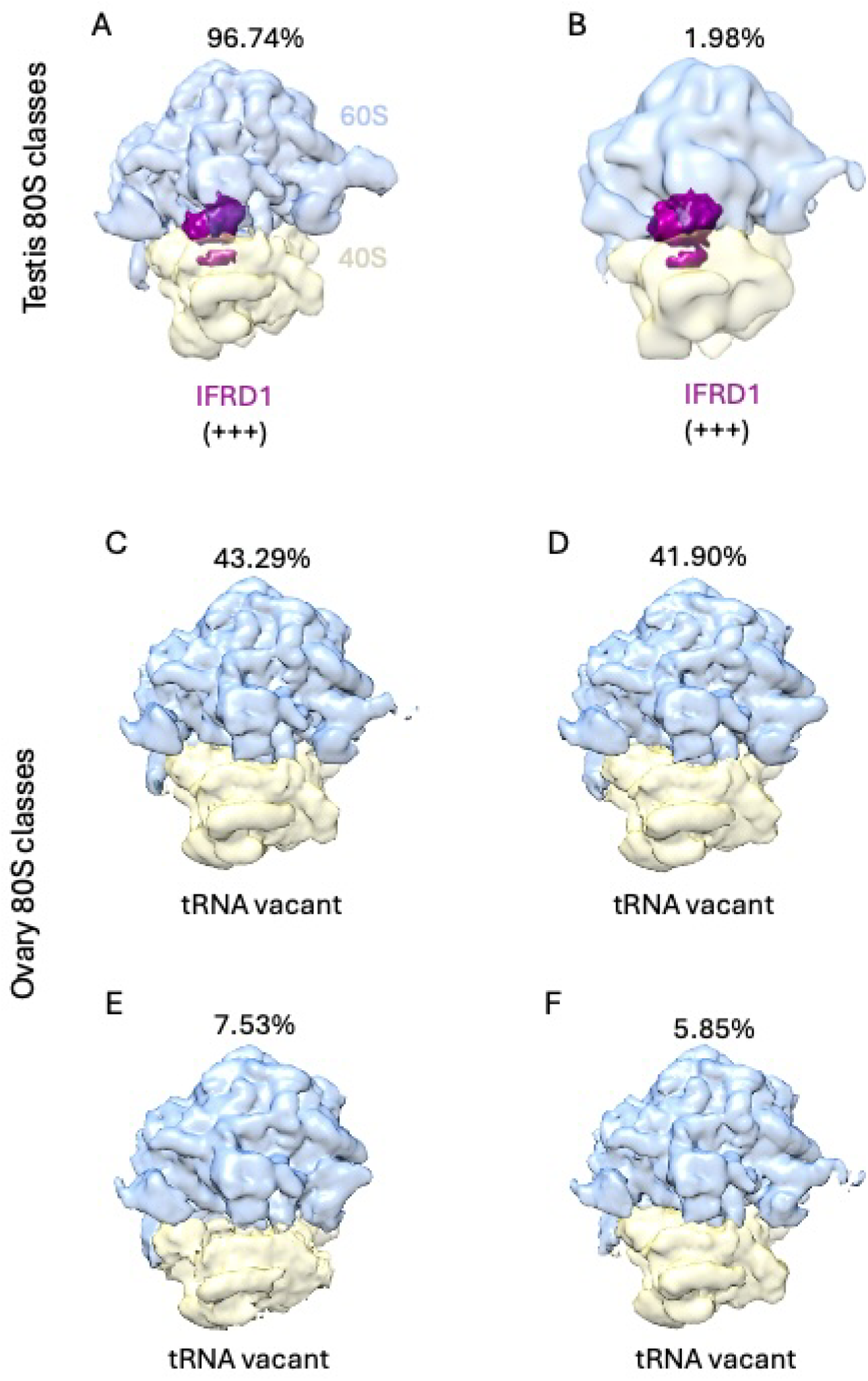
Testis and ovary 80S monosomes are not actively translating. Significant 3D classes (>1% total particles in dataset and >300 particles) of the testis 80S dataset (A-B) from ∼47,000 particles; and ovary 80S dataset (C-F) from ∼186,000 particles. (A and B) 2 classes of the testis 80S dataset, representing 96.74% and 1.98% of all testis 80S particles, contained a strong IFRD1 (purple) density in the mRNA channel. (C to F) All 4 significant classes from the ovary 80S dataset, representing 43.29%, 41.90%, 7.53% and 5.85%, lacked tRNA at the mRNA channel. Percentage of particles in each class is given above each class. IFRD1 position is indicated along with score of confidence with +++ representing high confidence.

### Ribosome complexes derived from polysomes are actively engaged in translation

As a positive control, we also performed mRNA channel focused classification on 2 samples expected to be highly active in translation: a) testis polysomes and b) 80S ribosomes derived from foot-printed embryo polysomes. These datasets contained ∼10,000 and ∼34,500 particles, respectively, and resulted in averages at 4.9 and 4.7 Å resolution. Over 97% of testis polysomes were tRNA occupied and therefore engaged in translation, exhibiting tRNA in the P/P and A/A sites (Figure S3A). Similarly, RNaseI footprinted polysomes from 0-2 hr embryos resulted in 4 significant classes, all of which were tRNA occupied and included P/E and A/P sites, E/E and P/P sites, and P/P and A/A sites (Figures S3B-E).

Overall, this analysis revealed that there is a clear contrast in the translational activity of 80S monosomes across different tissues (Figure 3, Supplemental Table 1). Head and embryo 80S monosomes are highly translationally active, testis 80S monosomes are inhibited and ovary 80S monosomes are vacant. As expected polysomal ribosomes are actively translating.

**Figure 3:**
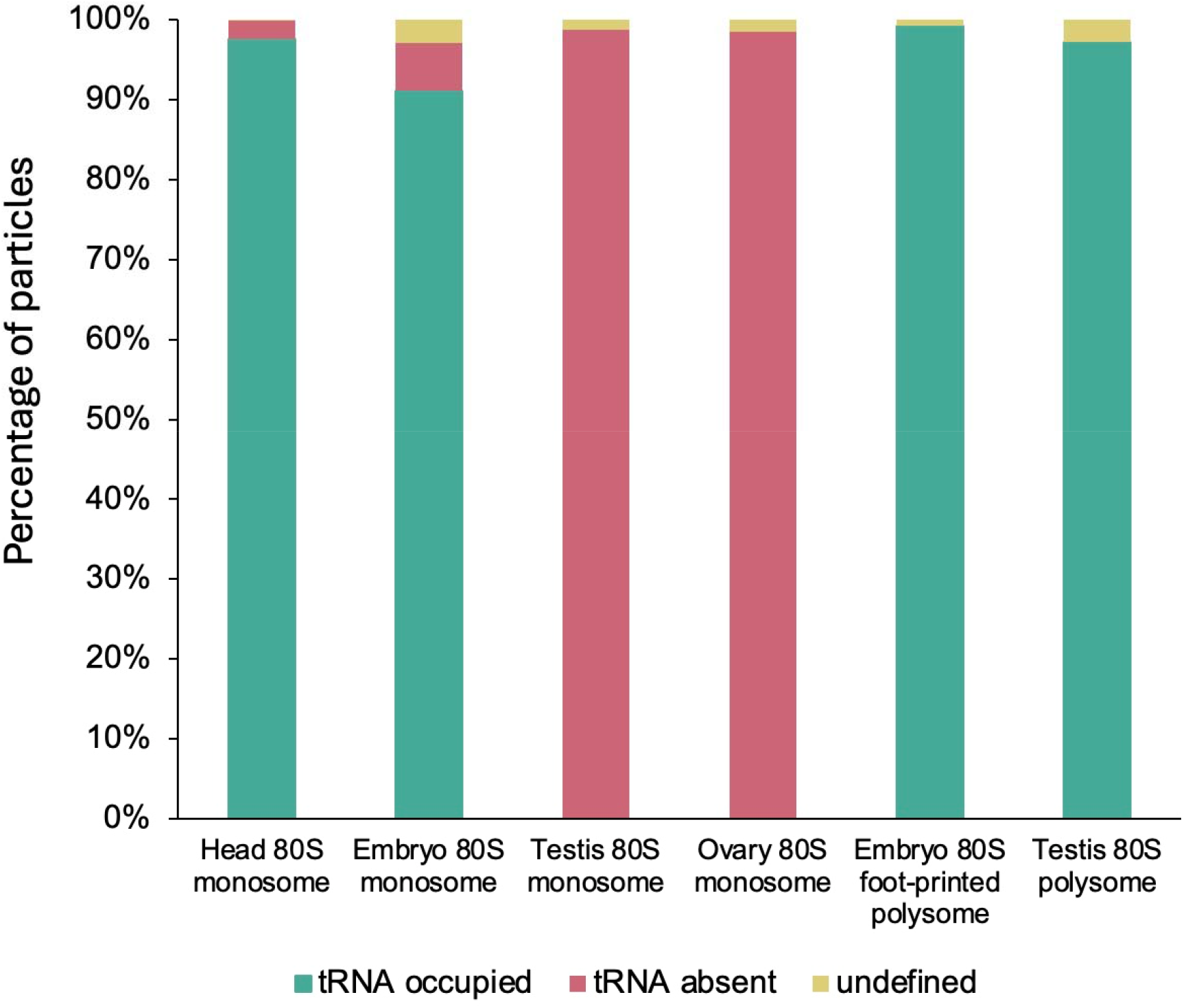
tRNA occupancy levels vary substantially across tissues. Bar chart showing percentages of particles for each dataset which either have tRNA densities present, absent, or resolution prevents definition (undefined).

### Translating 80S monosomes are stable in high salt conditions

To characterise the 80S monosomes in more detail we assessed their sensitivity to high KCl. Most ribosome purification buffers contain 0-100 mM KCl (Aspden and Jackson 2010; Sinha et al. 2020) but high KCl (e.g. 200mM) has been shown to dissociate vacant ribosomes not engaged with mRNAs (Martin and Hartwell 1970; Infante and Baierlein 1971; Saba et al. 2023). Therefore, to biochemically confirm that most 80S monosomes from heads are translating and therefore bound to mRNAs, we treated ovary 80S monosomes and head 80S monosomes with increasing concentrations of KCl (0, 100 and 200 mM) and monitored the effects of the incubation using sucrose gradients (Figure S4). At 0 mM and 100 mM KCl, which are widely used in sucrose gradient analysis, both ovary and head 80S remained associated (Figure S4, 0 and 100 mM KCl). At 200 mM KCl head ribosomes remained as 80S monosomes (Figures S4A and B, 200 mM), whilst the majority (∼90%) of ovary 80S monosomes were disassociated into 40S and 60S subunits (Figures S4C and D, 200 mM). The observation that head 80S monosomes are mostly insensitive but the majority of ovary 80S monosomes are sensitive, supports the conclusion that most head 80S monosomes are translating, whilst most ovary 80S monosomes are vacant.

### Head 80S monosomes and polysomes translate different mRNA pools

Given head 80S monosomes are engaged in active translation we sought to determine which mRNAs they translate and if they differ from the mRNAs translated by polysomes. RNA-Seq was performed followed by differential expression analysis on RNAs bound by 80S ribosomes or polysomes compared to total cytoplasmic RNA (Figure 4A). PCA showed that replicates from these different subcellular fractions clustered together but represented distinct RNA populations (Figure S5A). 784 genes were found to be preferentially translated by head 80S monosomes and 788 preferentially translated by head polysomes (Figure 4B). GO term analysis on these enriched mRNA populations revealed that mRNAs preferentially translated by monosomes were enriched for mRNAs encoding the translational machinery (Figure 4C). Whilst polysome-translated mRNAs were enriched for mRNAs encoding proteins involved with neural function (Figure 4D). This list of head monosome-enriched genes included 74 ribosomal proteins, 17 eIFs, 6 eEFs, RACK1, Poly A Binding Protein (PABP) and translationally controlled tumor protein (tctp). The mRNAs encoding these proteins have been previously shown to contain pyridimide-containing TOP motifs, which are highly conserved between human and *Drosophila* (Meyuhas and Dreazen 2009). Therefore, we analysed the 5’-UTR sequences of the 80S monosome enriched mRNAs, searching for over-represented motifs. The most enriched motif, found in 15% of these mRNAs was the TOP motif (Figures 4E and S5B). By analysing the attributes of the mRNAs preferentially translated by head 80S monosomes and polysomes, we discovered that monosome-translated mRNAs had significantly shorter ORFs (Figure 4F), shorter 5’-UTRs (Figure 4G) and shorter 3’-UTRs (Figure S5C). Monosome-translated mRNAs were also more likely to contain uORFs compared to polysomes-translated mRNAs (one-sided Fisher Exact Test p-value = 0.001645). Overall, these results show that head 80S and head polysomes translate different populations of mRNAs.

**Figure 4:**
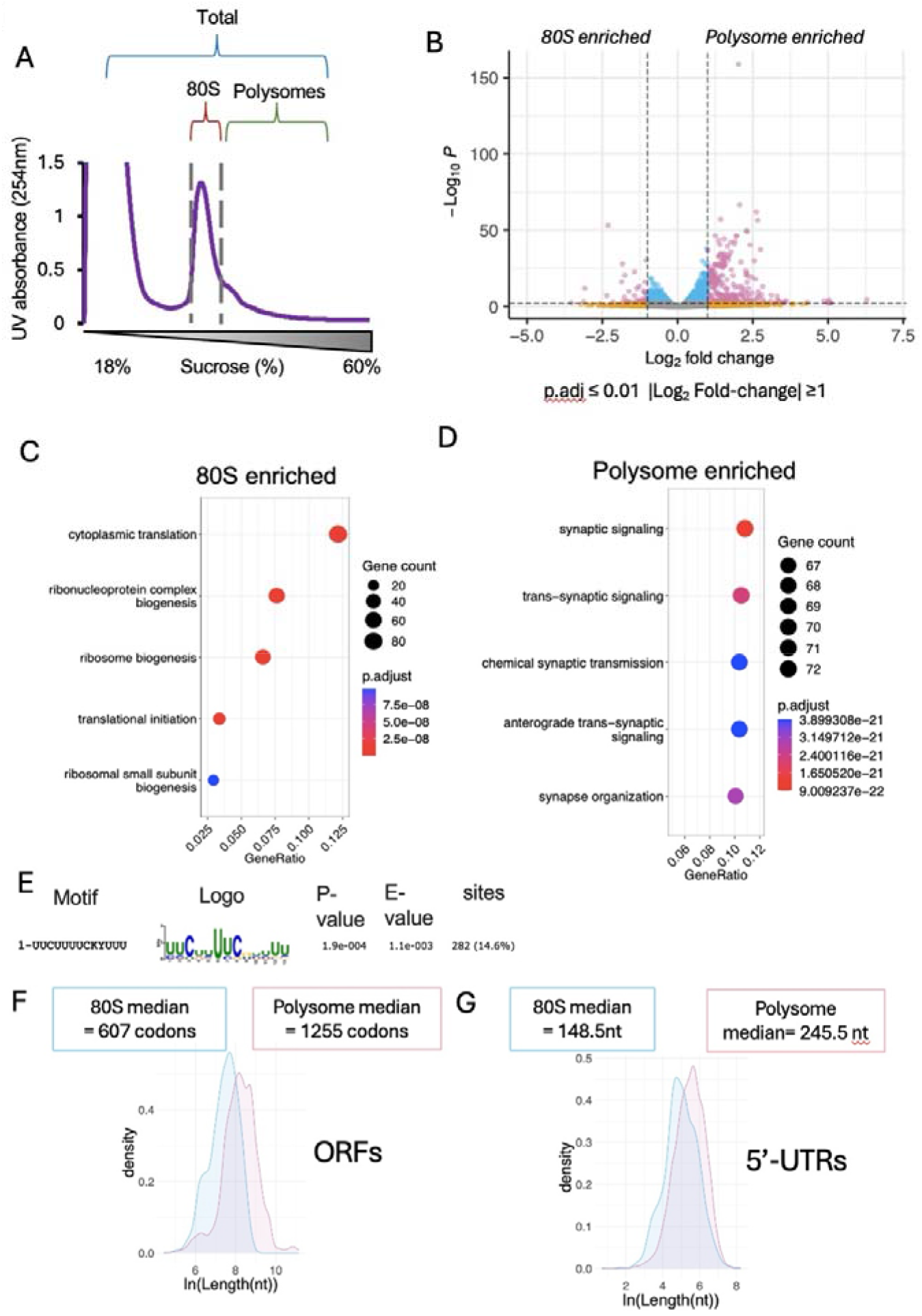
Head 80S monosomes and polysomes translate distinct mRNA populations. (A) Schematic of RNA-Seq samples from head ribosomal complexes. (B) Volcano plot of differentially enriched mRNAs between 80S monosomes and polysomes. Over-represented GO terms found in (C) 80S monosomes enriched mRNAs and (D) polysome-enriched mRNAs. (E) Most enriched motif in the 5’-UTRs of 80S monosome enriched mRNAs. Distributions of (F) ORF length (one-sided KS Test p-value = 4.5e^-167^) and (G) 5’-UTR length (one-sided KS Test p-value = 7.2e^-56^)(ln of length in nt) for 80S monosome and polysome enriched transcripts, with medians provided (80S monosome; blue, polysome; pink).

## Discussion

The view that 80S monosomes represent lowly translating ribosomal complexes is mostly derived from tissue cultured cells. However, this has not been fully characterised *in vivo*. To test this, we purified and characterised 80S monosomes and polysomes from different *D. melanogaster* tissues. Our cryo-EM analysis revealed substantial differences exist in the translational status of these 80S monosomes. The vast majority of 80S monosomes in the head (∼98%) and embryo (∼91%) contain tRNAs, indicating they are actively translating. Whilst 80S monosomes in the testis are inactive by way of IFRD1 binding within the mRNA channel. This is in contrast to ovary 80S monosomes, which are vacant and can be disassociated with high KCl (200 mM). As expected, 80S particles from polysomes in both embryo and testis are occupied with tRNAs, therefore actively translating.

*Danio rerio* and *Xenopus laevis* eggs and embryos, have been found to be bound byseveral dormancy factors (Leesch et al. 2023). Dap1b-eIF5a and Habp4-eEF2 act to stabilise 80S monosomes in eggs, repressing translation, ready for activation after fertilisation (Leesch et al. 2023). This is similar to our testis 80S monosomes, which have IFRD1 blocking the mRNA channel, and to IFRD2-inhibited ribosomes from rabbit reticulocyte lysate (Brown et al. 2018). Whilst our *Drosophila* embryos (0-2 hr) represent a stage where translational repression has been relieved, similar to 3 hours post fertilisation in (Leesch et al. 2023). Therefore, we did not find evidence of any 80S monosome dormancy in 0-2 hr *Drosophila* embryos, similar to previous cryo-EM of *D. melanogaster* 0-16 hr (Gebauer et al. 1999) embryo 80S monosomes, which showed tRNA occupancy and no dormancy factors (!!! INVALID CITATION !!! (Anger et al. 2013)).

The fact that 80S monosomes in embryos are the majority of the actively translating ribosomal complexes is consistent with the high levels of translation of uORFs in 0-2 hr embryos detected by Ribo-Seq (Arava et al. 2003). Given the small size of uORFs and therefore limited space for ribosomes to bind, only single 80S are expected to be translating at any one time. During early embryogenesis when transcription is silent, translational control is essential and uORF translation is a widespread mechanism of translational repression of main ORFs in maternal mRNAs at this time (Dunn et al. 2013).

To determine the translational contribution of 80S monosomes in the head compared to polysomes we performed RNA-Seq. This showed that these two types of translational complexes have different translational preferences. Specifically, head polysomes preferentially translate mRNAs with neuronal functions. Whilst head 80S monosomes preferentially translate mRNAs encoding the translational machinery itself. These 80S monosome translated mRNAs contain TOP motifs, have short 5’-UTRs, contain short ORFs and are enriched for uORFs. Again, given that there is limited room on short ORFs and uORFs for ribosomes, our result fits with this restriction (Arava et al. 2003). This is also consistent with ribosome profiling of monosomes in *Saccharomyces cerevisiae* where most monosomes are actively elongating, and that monosomes translate short ORFs, uORFs, NMD targets and low-abundance regulatory proteins (Heyer and Moore 2016).

Head 80S monosome-translated mRNAs also tend to have shorter 5’-UTRs and 3’-UTRs not just compared to polysome-translated but also shorter than average in invertebrates (Mignone et al. 2002). The translational regulation of 5’TOP mRNAs has been shown to involve a shift in their translation between polysomes and monosomes (Schneider et al. 2022). In response to stresses such as starvation in tissue culture cells, the RNA-binding protein LARP1 represses 5’TOP mRNA translation, but basal levels of translation continue to occur by 80S monosomes. Our heads come from well fed *D. melanogaster* so starvation is unlikely to be the signal for our ribosomes, but a comparable regulatory network may exist in the head.

Our evidence for translation of neuronal function mRNAs by head polysomes may seem in contrast to previous evidence indicating 80S monosomes actively translate synaptic mRNAs in neuronal processes (Biever et al. 2020). However, we have isolated ribosomes from the whole fly head, only a proportion of which is brain. Therefore, our 80 monosomes could come from a variety of cell types in addition to neurons, such as glial cells, fat bodies and muscles. In fact, Ribo-Seq in *Drosophila* glia has revealed a substantial amount of uORF translation (Ichinose et al. 2023), which is likely contributing to 80S monosome translation in our analysis. Comparison of Ribo-Seq from glial cells and neurons indicates translational control is substantially different, and our analysis of whole heads will contain both types of translation events, along with the multiple other cell types, making direct comparison with neurons alone complex (Biever et al. 2020).

Overall, our results reveal that the translational status of 80S monosomes varies substantially across tissues, where these ribosomes represent the dominant translational complex compared to tissue culture cells.

## Methods

### Fly husbandry and stocks

Flies were kept in a 25°C humidified room with a 12:12 hour light:dark cycle and raised on standard sugar-yeast-agar medium in 6 oz Square Bottom Bottles (Flystuff).

### Tissue harvest for cryo-EM and RNA-Seq

5 mL of 0-5 day old whole flies were snap frozen in liquid nitrogen (LN_2_) and subjected to mechanical shock to detach heads. Heads were isolated by passing through a 1 mm mesh filter with LN_2_. 1.5 g of embryos were collected at 0-2 hr from laying plates [300 mL grape juice concentrate (Young’s Brew), 25 g agar, 550 mL dH2O, 10% nipagin]. Laying plates were scored and applied with yeast paste consisting of active dried yeast (DCL), plates were placed in cages after pre-clearing for 2 hr. Embryos were transferred to 70 mm mesh filter, washed with dH_2_O, dried and flash frozen in LN_2_. 500-1000 pairs of testes were harvested from 1-to 4-day-old males in 1× PBS with 2 mM DTT and 1 U/μl RNAsin Plus and flash frozen in LN_2_ (Hopes et al. 2022). ∼300 pairs of ovaries were dissected from 3-6 day old females in 1× PBS (Lonza) with 1 mM DTT (Sigma) and 1 U/μl RNAsin Plus (Promega) and flash frozen in LN_2_ (Hopes et al. 2022).

### Ribosome purification for cryo-EM and RNA-seq

All stages of ribosome purification were performed on wet ice or at 4°C, wherever possible. Heads were transferred to an 8 mL glass Dounce with 1.5 mL lysis buffer A [10 mM Tris-HCl pH 7.5, 150 mM NaCl, 10 mM MgCl_2_ (Fluka), 1% IGEPAL CA-630 (Sigma), 1% Triton X-100, 0.5% sodium deoxycholate (Sigma), 2 mM DTT, 200 µg/ml cycloheximide, 2 U/mL Turbo DNAse (Thermo Fisher), 40 U/mL RNAsin Plus, 1× EDTA-free protease inhibitor cocktail (Roche), 0.5% DOC] per gradient. Embryos were ground with a pre-chilled pestle and mortar in LN_2_ and incubated with 1.5 mL lysis buffer A per gradient. Ovaries and testes were ground using RNase-free 1.5 mL pestles (SLS) in 500 mL lysis buffer B per gradient [50 mM Tris-HCl pH 8 (Sigma), 150 mM NaCl, 10 mM MgCl_2_, 1% IGEPAL CA-630, 1 mM DTT, 100 µg/Ml cycloheximide, 2 U/µL Turbo DNase, 0.2 U/µL RNasin Plus, 1× EDTA-free protease inhibitor cocktail] (Hopes et al. 2022). All samples were incubated for 30 minutes with occasional agitation. To obtain the cytoplasmic lysate, both head and embryo preparations were centrifuged to remove cell debris and floating fat at 17,000 × g for 10 min.

Cytoplasmic lysates were loaded onto a 18-60% (w/v) sucrose gradient [50 mM Tris-HCl pH 8.0, 150 mM NaCl, 10 mM MgCl_2_, 100 mg/ml cycloheximide, 1 mM DTT, 1× EDTA-free protease inhibitor cocktail (Roche)] and ultra-centrifuged for 3.5 hours at 170,920 × g at 4°C. Fractions were collected using a Gradient StationBiocomp equipped with a fraction collector (Gilson) and Econo UV monitor (BioRad). Fractions corresponding to either monosomes or polysomes were combined. These fractions were concentrated using 30 kDa column (Amicon Ultra-4 or Ultra-15) at 4°C and buffer exchanged (50 mM Tris–HCl pH 8, 150 mM NaCl, 10 mM MgCl_2_) until final sucrose ≤0.1%. The foot-printed 80S polysomes were isolated from pooled polysomal fractions of the embryo extract ribosome purification sucrose gradient and treated with RNaseI (4 U/area under the curve; AUC at 254nm at 4°C overnight. SuperRNAsin was added for 5 minutes at 4°C, preventing over digestion of mRNA. A second ribosome purification step was carried out on this sample, isolating the 80S fractions. Fractions were pooled and diluted to <0.1% sucrose and concentrated using Amicon Ultra-4 centrifugal filter units (MWCO 30 kDa) to >200 nM. Samples were quantified using Qubit Protein Assay Kit (Invitrogen) (Hopes et al. 2022).

### Application to cryo-EM grids

Purified ribosomes were diluted as required with dilution buffer (50 mM Tris-HCl pH 8.0, 150 mM NaCl, 10 mM MgCl_2_). Copper grids covered with a lacey carbon and an ultrathin layer of carbon (Agar Scientific) were glow discharged for 30 seconds (easiGlow, Ted Pella) and 3 µL of purified ribosomes were added to the grid surface in chamber at 4°C and 95% humidity conditions. Sample excess was blotted and grids were vitrified by plunge-freeing in liquid ethane cooled by liquid nitrogen using the EM GP plunge freezer (Leica)(Hopes et al. 2022).

### Cryo-EM data collection

For all samples, cryo-EM data collection was carried out using FEI Titan Krios (Thermo Fisher) transmission electron microscope with an accelerating voltage of 300 keV. Data was recorded on a Falcon III direct electron detector in integrating mode at a pixel size of 1.065 Å. The total number of collected micrographs for the head 80S monosomes was 14,827, while for the foot-printed testis polysomes, a total of 8,128 micrographs were collected. For both samples a total electron dose of 82 e^-^/Å^2^ partitioned into a dose of 1.37 e^-^/Å^2^ per fraction (60 fractions) was employed. Data collection for all other samples have been described previously (Hopes et al. 2022).

### Cryo-EM image processing

Cryo-EM image processing (Thompson et al. 2019; Zivanov et al. 2018). Motion correction and CTF estimation were performed using MOTIONCORR and gCTF, respectively (Zhang 2016; Li et al. 2013). For the head 80S monosomes 735,663 particles were picked using crYOLO’s general model (Wagner et al. 2019). For the testis foot-printed polysomes, autopicking was performed in Relion, resulting in 185,095 particles. Relion was used for further processing. 2 rounds of reference free 2D classification were used to align and classify autopicked particles into 200 classes. 2D classifications were used to iteratively remove “junk” particles selected from autopicking. After both rounds of 2D classification, the head 80S dataset resulted in 610,605 particles while the embryo footprinted polysomes dataset resulted in 69,712 particles. The previously produced *D. melanogaster* testis 80S ribosome average was used as a reference average for 3D classification (Hopes et al. 2022). These particles were further 3D classified, refined and postprocessed, resulting in averages at a final resolution of 3.0 Å for the heads 80S dataset (Figure S2) and of 4.7 Å for the embryo footprinted polysomes.

### Focussed classification

The proportion of 80S monosomes and polysomes engaged in active translation in the *D. melanogaster* head, embryo, testis and ovary tissues was determined by tRNA occupation assessment by focused classification of the different datasets. The head 80S monosome dataset contained 610,605 particles (as above), the embryo 80S monosome dataset 11,446 particles, the embryo foot-printed 80S polysome dataset 34,603 particles, the testis 80S monosome dataset 46,878 particles, the testis 80S polysome dataset 10,392 particles and the ovary 80S monosome dataset 185,913 particles. After 3D refinement carried out using unbinned particles, as described above, particles were then binned 5 times to expedite image processing (5.325 Å pixel size), given that tRNAs are easily distinguished at the theoretical maximum resolution achievable (10.65 Å).

A mask around the mRNA channel was then generated, and a 10-class 3D classification without alignment was performed. To resolve the full 80S monosome structure whilst maintaining the structural detail from the masked focused classification, a final unmasked reconstruction of each class was performed. This allowed imaging the full 80S monosome, aiding orientating and assessing the mRNA channel and tRNA positions in the context of the whole 80S monosome.

### tRNA and IFRD1 atomic model fitting

To determine the conformation of tRNAs in the *D. melanogaster* tissue ribosomes, reference eukaryotic tRNA atomic models were chosen from variety of organisms, as only the P/P and E/E tRNAs are currently available from *D. melanogaster* ribosome atomic models. These included the A/A tRNA from *S. cerevisiae* (PDB 5GAK), the A/P and P/E tRNAs from rabbit reticulocyte lysate (PDB 6HCJ) and P/P and /E tRNAs from *D. melanogaster* (respectively, PDB 6XU7 and 4V6W). The IFRD1 atomic model was selected from the *D. melanogaster* testis 80S monosome atomic model (PDB 6XU6) (Hopes et al. 2022).

### RNA-Seq

Head tissue was harvested, and ribosomes purified as above (Hopes et al., 2022). 10% of cell lysate was retained for sequencing total RNA. Fractions were isopropanol-precipitated (1 vol isopropanol, 300 mM NaCl final) overnight at -80°C, then centrifuged at 17,000 x g for 20 min and washed in 70% ethanol. 80S monosome fractions were combined, as were polysome fractions. Total RNA and sucrose gradient fractions were treated with TURBO DNase (Ambion) and purified using Quick-RNA Microprep Kit (Zymo). Samples were rRNA depleted with QIAseq FastSelect –rRNA Fly Kit (QIAGEN). Sequencing libraries were prepared with the TruSeq Stranded mRNA kit (Illumina) and run on the NextSeq 2000, 100 cycle lane (Illumina).

### RNA-Seq analysis

RNA single-read sequencing quality control was assessed through FastQC (www.bioinformatics.babraham.ac.uk/projects/fastqc) and multiQC (Ewels et al. 2016). An average of 45 million reads were sequenced per sample. Both adapters and low-quality bases (QV < 20) were trimmed from reads’ extremities using Trimmomatic (Bolger, Lohse, and Usadel 2014) with a minimum read length of 25 bp. All libraries were mapped against the *D. melanogaster* BDGP6.32 release-109 reference genome (http://ftp.ensembl.org/pub/release-09/fasta/drosophila_melanogaster/dna/Drosophila_melanogaster.BDGP6.32.dna.toplevel.fa.gz) through STAR aligner (Dobin 2013) with default parameters. STAR-generated sorted BAM output files were used for assigning read counts to gene features with featureCounts (Liao, Smyth, and Shi 2014) with the following parameters: -M -O --fraction -s 2. Drosophila_melanogaster.BDGP6.32.109.gtf annotation file from ensembl was used.

Read counts table generated by featureCounts was then used as input for differential expression (DE) analyses relying on the DeSeq2 negative binomial distribution model through a local fitting type and a 0.05 FDR threshold (Love 2014). DE pairwise comparisons were performed as 80S-vs-Polysomes, Total-vs-80S, and Total-vs-Polysomes, being the second term the numerator in the fold-change ratio. EnhancedVolcano (https://bioconductor.org/packages/release/bioc/html/EnhancedVolcano.html) was employed for an overall DE visualization through volcano plots. GO- and KEGG-based gene enrichment analyses for individual DEGs lists were performed by clusterProfiler Bioconductor package (Wu et al. 2021) under R v4.1.0, setting adjusted p-value < 0.05. Both 5’- and 3’-UTRs were extracted from Drosophila_melanogaster.BDGP6.32.109.gtf. An empirical cumulative distribution function (ECDF) plus Kolmogorov-Smirnov Tests were run within the R v4.1.0 environment to assign significance to UTR lengths comparison between conditional groups (80S, Polysomes, and Total RNA). STREME from the MEME suite v5.4.1 (Bailey et al. 2015) was employed for motif screening within the UTRs of both 80S- and Polysome-enriched transcripts under the following parameters: --rna --minw 5 --maxw 15 --thresh 0.01. A negative control set of randomly-selected non-DEGs’ UTRs in the same amount of the DEGs’ ones was used for the --n option.

### KCl treatment

For heads, 1-5 day-old flies were sedated using ice. On an iced glass petridish, 50 heads were removed and placed in 173 µL dissecting buffer (Schneider’s media with x1 EDTA-free protease inhibitor cocktail (Roche), x1 RNasin Plus (Promega) and 100 µg/mL cycloheximide). This was repeated to 150 heads per gradient. For ovaries, 1-5 day-old flies were sedated using ice. On a dissecting dish, 20 ovaries were removed and placed in 173 µL dissecting buffer. Heads and ovaries were ground using RNase-free 1.5 mL pestles (SLS). 207 µL lysis buffer A plus 100 or 200 mM KCl. Samples were lysed on ice for ≥30 mins, then centrifuged at 17,000 x g for 10 mins at 4°C.

Sucrose gradients were made from 2 mL each of 50/47/42/34%/26/18% sucrose, 50 mM Tris-KCl pH 8.0, 150 mM NaCl, 10 mM MgCl_2_, 100 mg/ml cycloheximide, 1 mM DTT, 1× EDTA-free Protease Inhibitor cocktail (Roche), and 0, 100 mM or 200 mM KCl). Cytoplasmic lysates were loaded onto a 18-50% (w/v) sucrose gradient and ultra-centrifuged in an SW40Ti rotor (Beckman) for 4:15 hours at 170,920 × g at 4°C and fractionated as above.

## Funding

AB and MA were funded from BBSRC DTP, BB/M011151/1. AT was funded by Wellcome Trust 102174/B/13/Z. TH, JF and JA were funded by BBSRC grant BB/S007407/1 and BBSRC BB/X003086/1. BF was funded by BB/X003086/1.

## Competing Interest Statement

Authors have no competing interests.

Acknowledgments

We thank the Astbury Biostructure Laboratory (ABSL) Facility Staff from the Electron Microscopy Facility for assisting with cryo-EM data collection. We thank the Next Generation Sequencing facility, at St James University Hospital, Leeds, UK for performing Next Generation Sequencing. Parts of this work were undertaken on ARC3, part of the High Performance Computing facilities at the University of Leeds, UK.

## Author contributions

AB performed experiments and analysed the data. TH conceived, designed and performed experiments for this study, and interpreted data. EV, AT, BF and MA performed experiments for this study and analysed the data. JF and JLA conceived the work, designed the experiments and interpreted the data. All authors contributed to manuscript writing, revision and have approved the submitted version.

## Notes

### Competing Interest Statement

The authors have declared no competing interest.

