## Supplemental for "Translational activity of 80S monosomes varies dramatically across different tissues"

### Slide 1
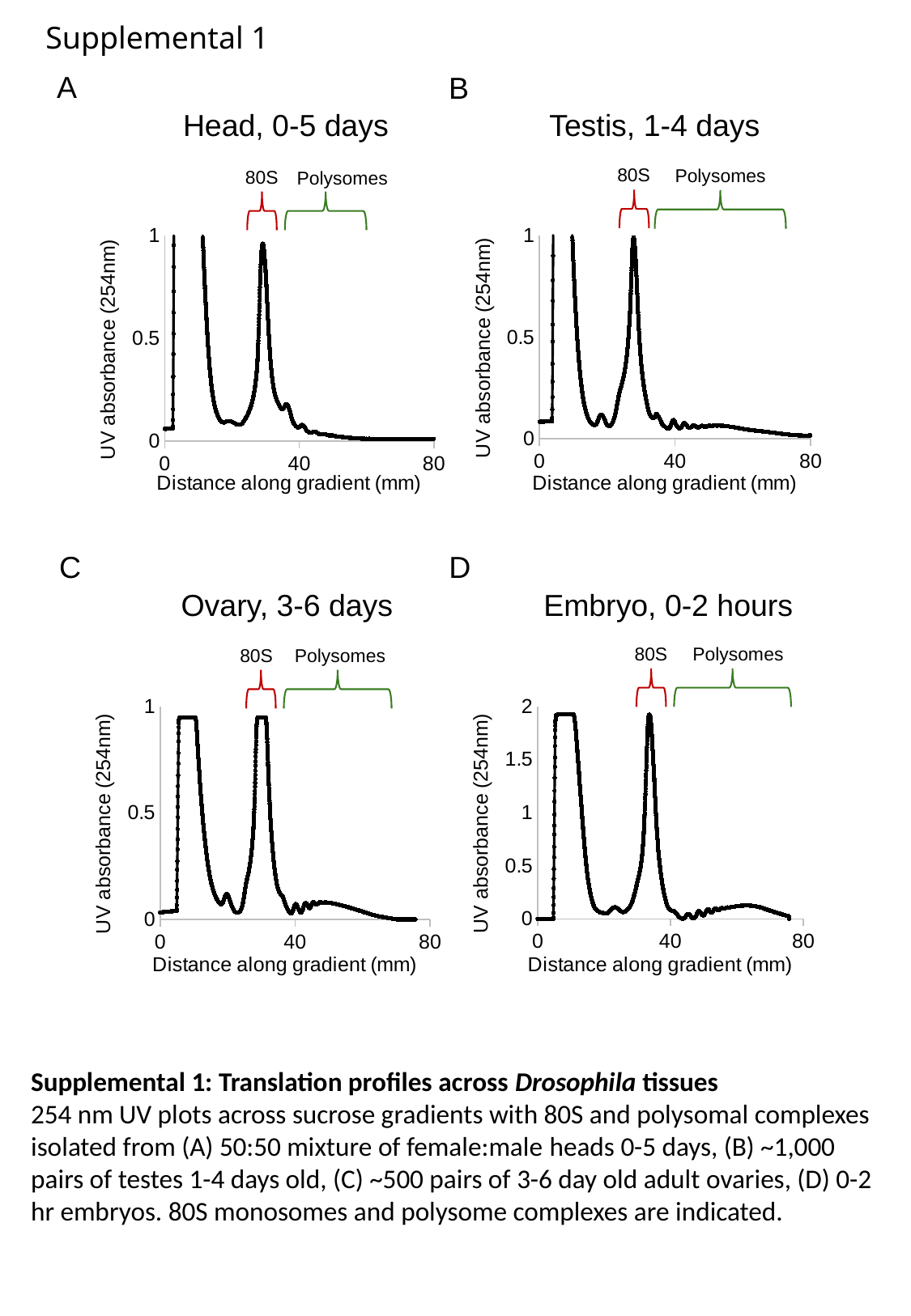

Supplemental 1
A
B
Testis, 1-4 days
Head, 0-5 days
80S
Polysomes
80S
Polysomes
#### Chart
| Category | TMT3_testies |
|---|---|
#### Chart
| Category | TMT3_heads_2 |
|---|---|C
D
Embryo, 0-2 hours
Ovary, 3-6 days
80S
Polysomes
Polysomes
80S
#### Chart
| Category | TMT1_ovaries |
|---|---|
#### Chart
| Category | TMT2_embryos |
|---|---|
Supplemental 1: Translation profiles across Drosophila tissues
254 nm UV plots across sucrose gradients with 80S and polysomal complexes isolated from (A) 50:50 mixture of female:male heads 0-5 days, (B) ~1,000 pairs of testes 1-4 days old, (C) ~500 pairs of 3-6 day old adult ovaries, (D) 0-2 hr embryos. 80S monosomes and polysome complexes are indicated.

### Slide 2
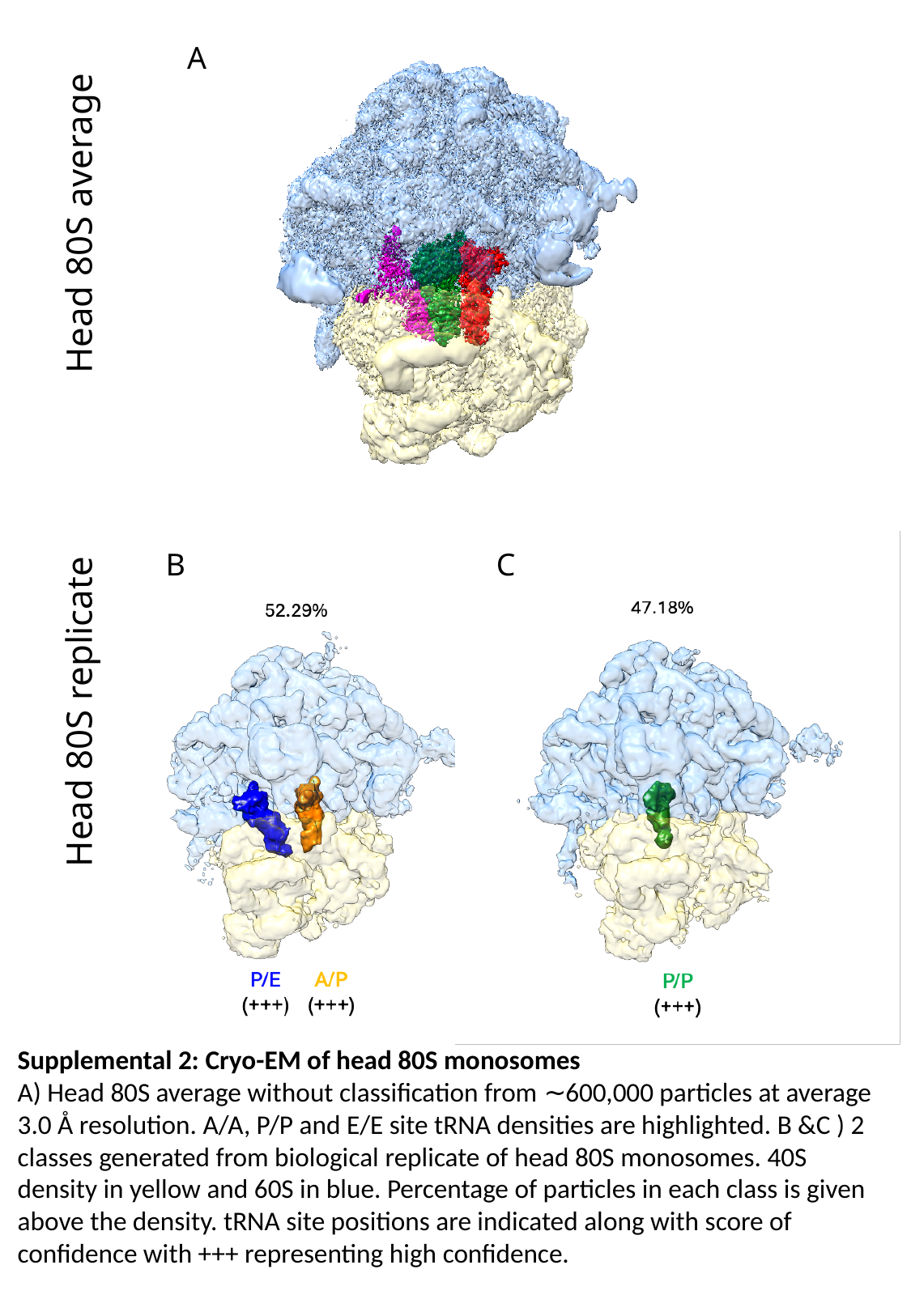

A
Head 80S average
B
C
Head 80S replicate
Supplemental 2: Cryo-EM of head 80S monosomes
A) Head 80S average without classification from ∼600,000 particles at average 3.0 Å resolution. A/A, P/P and E/E site tRNA densities are highlighted. B &C ) 2 classes generated from biological replicate of head 80S monosomes. 40S density in yellow and 60S in blue. Percentage of particles in each class is given above the density. tRNA site positions are indicated along with score of confidence with +++ representing high confidence.

### Slide 3
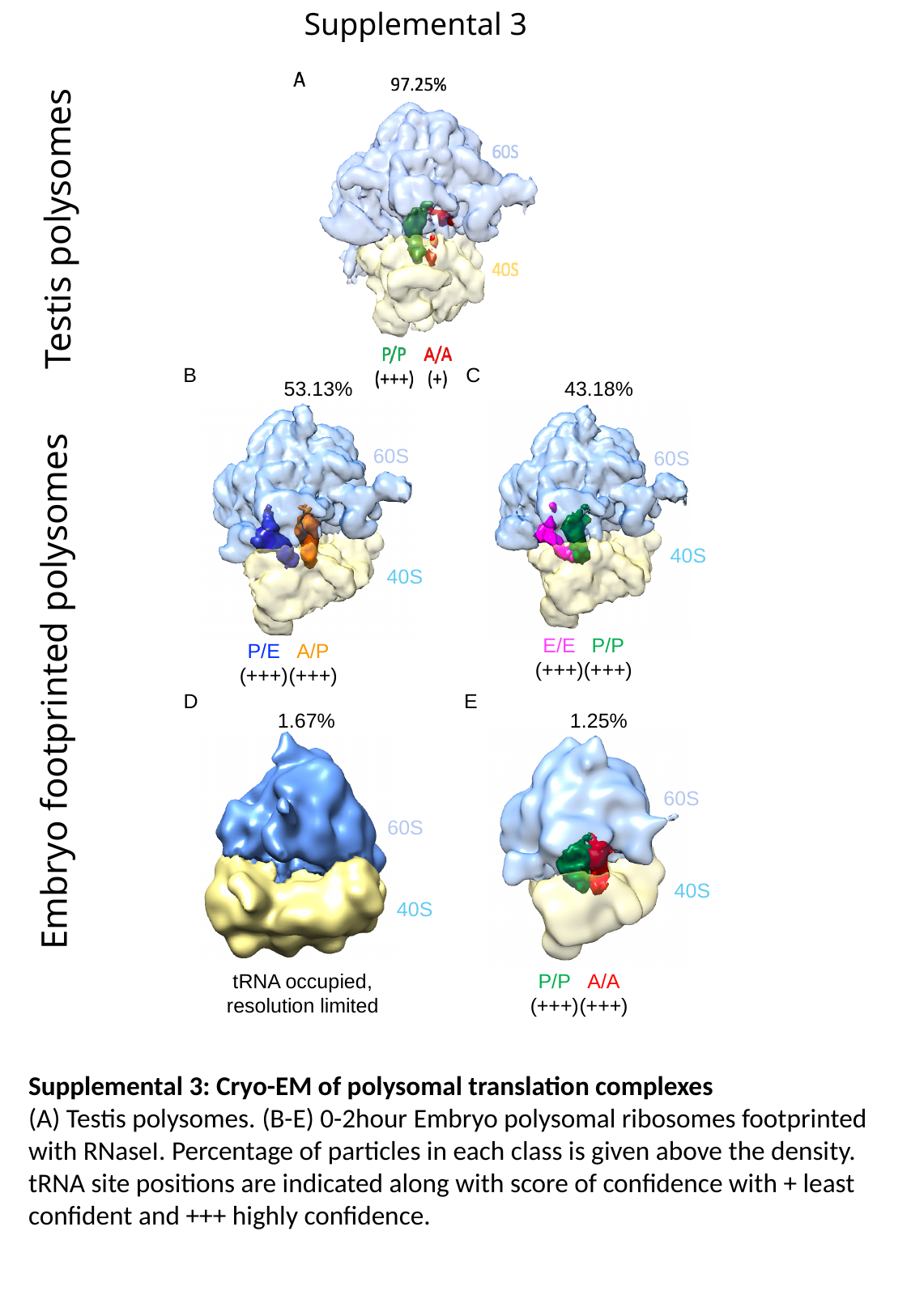

Supplemental 3
Testis polysomes
B
C
53.13%
43.18%
60S
60S
40S
40S
E/E
(+++)
P/P
(+++)
A/P
(+++)
P/E
(+++)
D
E
1.67%
1.25%
60S
60S
40S
40S
tRNA occupied, resolution limited
A/A
(+++)
P/P
(+++)
Embryo footprinted polysomes
Supplemental 3: Cryo-EM of polysomal translation complexes
(A) Testis polysomes. (B-E) 0-2hour Embryo polysomal ribosomes footprinted with RNaseI. Percentage of particles in each class is given above the density. tRNA site positions are indicated along with score of confidence with + least confident and +++ highly confidence.

### Slide 4
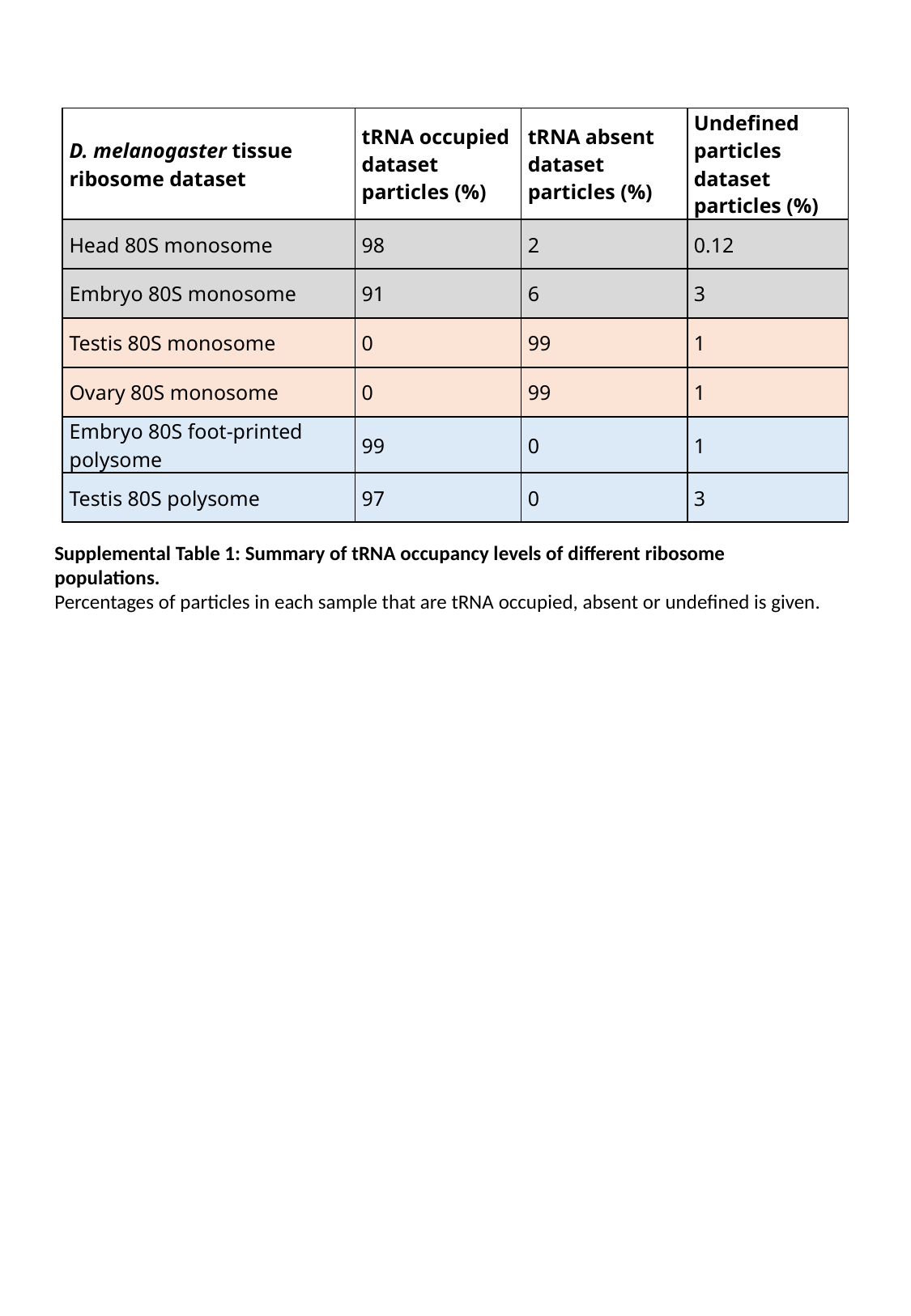

| D. melanogaster tissue ribosome dataset | tRNA occupied dataset particles (%) | tRNA absent dataset particles (%) | Undefined particles dataset particles (%) |
| --- | --- | --- | --- |
| Head 80S monosome | 98 | 2 | 0.12 |
| Embryo 80S monosome | 91 | 6 | 3 |
| Testis 80S monosome | 0 | 99 | 1 |
| Ovary 80S monosome | 0 | 99 | 1 |
| Embryo 80S foot-printed polysome | 99 | 0 | 1 |
| Testis 80S polysome | 97 | 0 | 3 |
Supplemental Table 1: Summary of tRNA occupancy levels of different ribosome populations.
Percentages of particles in each sample that are tRNA occupied, absent or undefined is given.

### Slide 5
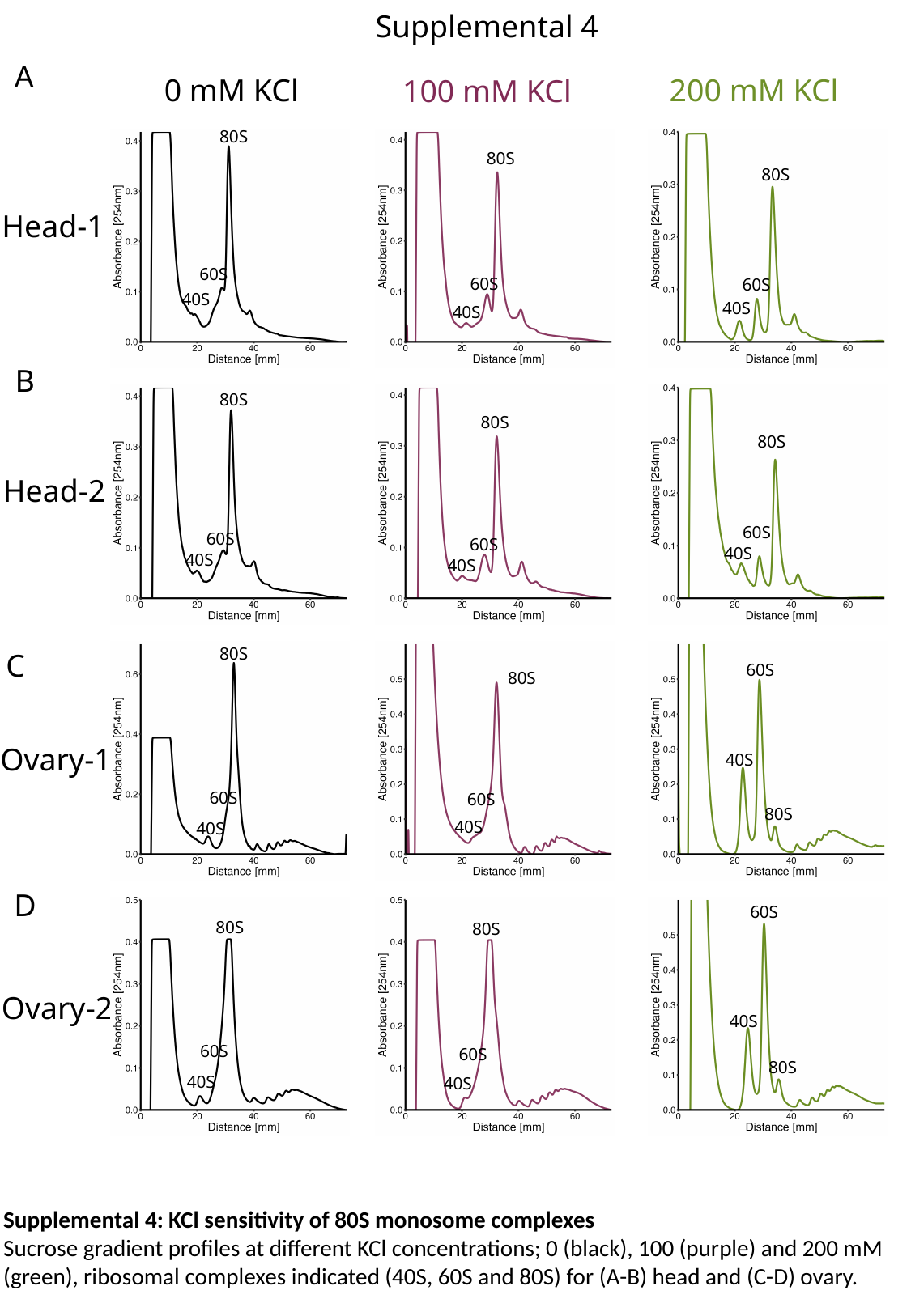

Supplemental 4
A
0 mM KCl
200 mM KCl
100 mM KCl
80S
80S
80S
Head-1
60S
60S
60S
40S
40S
40S
B
80S
80S
80S
Head-2
60S
60S
60S
40S
40S
40S
80S
C
60S
80S
Ovary-1
40S
60S
60S
80S
40S
40S
D
60S
80S
80S
Ovary-2
40S
60S
60S
80S
40S
40S
Supplemental 4: KCl sensitivity of 80S monosome complexes
Sucrose gradient profiles at different KCl concentrations; 0 (black), 100 (purple) and 200 mM (green), ribosomal complexes indicated (40S, 60S and 80S) for (A-B) head and (C-D) ovary.

### Slide 6
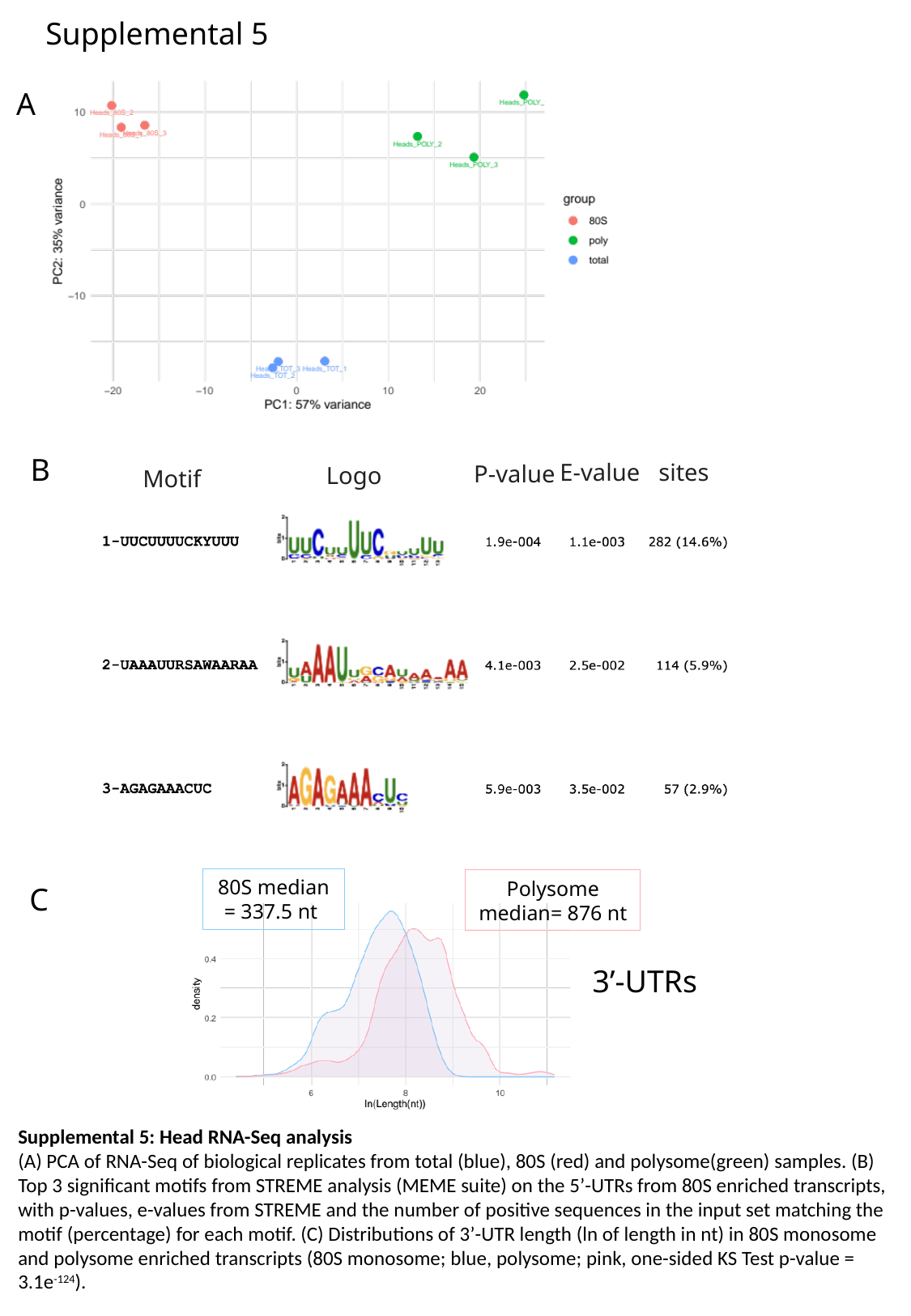

Supplemental 5
A
B
E-value
sites
P-value
Logo
Motif
80S median
= 337.5 nt
Polysome median= 876 nt
C
3’-UTRs
Supplemental 5: Head RNA-Seq analysis
(A) PCA of RNA-Seq of biological replicates from total (blue), 80S (red) and polysome(green) samples. (B) Top 3 significant motifs from STREME analysis (MEME suite) on the 5’-UTRs from 80S enriched transcripts, with p-values, e-values from STREME and the number of positive sequences in the input set matching the motif (percentage) for each motif. (C) Distributions of 3’-UTR length (ln of length in nt) in 80S monosome and polysome enriched transcripts (80S monosome; blue, polysome; pink, one-sided KS Test p-value = 3.1e^-124^).
